# Deciphering Transcriptional Regulation of Human Core Promoters

**DOI:** 10.1101/174904

**Authors:** Shira Weingarten-Gabbay, Ronit Nir, Shai Lubliner, Eilon Sharon, Yael Kalma, Adina Weinberger, Eran Segal

**Affiliations:** Department of Computer Science and Applied Mathematics, Weizmann Institute of Science, Rehovot 76100, Israel; Department of Molecular Cell Biology, Weizmann Institute of Science, Rehovot 76100, Israel; Department of Molecular Genetics, Weizmann Institute of Science, Rehovot 76100, Israel; Department of Genetics, Stanford University, Stanford, California, USA; Department of Biology, Stanford University, Stanford, California, USA; IVF Lab and Wolfe PGD-Stem Cell Lab, Racine IVF Unit, Lis Maternity Hospital, Tel-Aviv Sourasky Medical Center, Tel Aviv, Israel

## Abstract

Despite its pivotal role in regulating transcription, our understanding of core promoter function, architecture, and cis-regulatory elements is lacking. Here, we devised a highthroughput assay to quantify the activity of ∼15,000 fully designed core promoters that we integrated and expressed from a fixed location within the human genome. We find that core promoters drive transcription unidirectionally, and that sequences originating from promoters exhibit stronger activity than sequences originating from enhancers. Testing multiple combinations and distances of core promoter elements, we observe a positive effect of TATA and Initiator, a negative effect of BREu and BREd, and a 10bp periodicity in the optimal distance between the TATA and the Initiator. By comprehensively screening TF binding-sites, we show that site orientation has little effect, that the effect of binding site number on expression is factor-specific, and that there is a striking agreement between the effect of binding site multiplicity in our assay and the tendency of the TF to appear in homotypic clusters throughout the genome. Overall, our results systematically assay the elements that drive expression in core- and proximal-promoter regions and shed light on organization principles of regulatory regions in the human genome.

## INTRODUCTION

In contrast to the significant progress made in identifying the DNA elements involved in transcriptional regulation, our understanding of the rules that govern this process, namely how the arrangement and combination of elements affect expression, remains mostly unknown(*1, 2*). Advances in DNA synthesis and sequencing technologies have led researchers to tackle these questions using high-throughput approaches, yet most studies focus on enhancers. Traditionally, the core promoter region was viewed as a universal stretch of DNA that directs the pre-initiation complex to initiate transcription. However, core promoters are structurally and functionally diverse regulatory sequences. Hence, in the analysis of gene expression, it is necessary to understand and to incorporate the specific components of the core promoter(*3, 4*). However, our understanding of the core promoter function, architecture and cis-regulatory sequences is still lacking. With the growing appreciation of the importance of core promoters in determining gene expression, two recent studies measured the autonomous promoter activity of random sequences genome-wide in human and *Drosophila*(*5–7*). However, native promoters differ in many sequence elements making it hard to attribute the measured expression differences to any single sequence change. Thus, to infer the cis-regulatory elements governing core promoter activity, a large number of designed sequences in which specific elements are systematically varied in a highly controlled setting should be assayed.

To address fundamental questions in gene expression using fully designed sequences, we and others have developed massively parallel reporter assays probing the expression of various regulatory regions(*8–12*). However, since these measurements are performed using episomal plasmids they are limited in their ability to mimic the genomic context. Progress in this direction was recently made by integrating the reporter construct into the human genome using lentiviruses(*13–15*). However, lentivirus-mediated integration occurs in random locations along the genome and is thus susceptible to the effects of local chromatin environment and interaction with neighboring enhancers. The latter is of high importance to the measurements of core promoters due to core-promoter-enhancer specificity resulting in variability in core promoter activity when placed near different sets of enhancers (*16*).

Here, we present a new high-throughput method for accurately measuring ∼15,000 fully designed sequences from a fixed and predefined locus in the human genome. Using this system, we set to decipher the sequence determinants of core promoters and the proximal promoter region from broad aspects of mapping their location and orientation in the genome to in-depth characterization of the cis-regulatory elements driving their expression including core promoter elements and TF binding-site. To this end, we designed oligonucleotides, 200 basepairs in length, representing native and synthetic sequences. We assayed hundreds of genomic regions bound by the pre-initiation complex (PIC) in cells as well as thousands core promoters of endogenous genes. To systematically interrogate key sequences in core promoters, we designed oligos in which we tested all possible combinations and distances of the six common core promoter elements including the TATA-box, Initiator (Inr), upstream and downstream TFIIB recognition elements (BREu and BREd), motif ten element (MTE), and downstream core promoter element (DPE). In addition, we performed TF activity screen for 133 binding-sites and designed ∼1000 promoters to dissect the effect of homotypic sites number on expression.

Our results uncovered a positive relationship between the binding intensity of the pre-initiation complex in cells and the activity of core promoters. We show that although core promoters are present in both promoters and enhancers they are more active in promoter regions and generally drive unidirectional transcription. Our analysis of core promoter elements demonstrates a positive effect on expression of the TATA and the Inr elements and a negative effect of the BRE upstream and downstream elements. In addition, we find that the distance between the TATA and the Inr has an effect on expression with higher activity at distances of 10, 20 and 30bp, matching the ∼10bp periodicity of the DNA double helix. Finally, we show that the effect of binding site multiplicity on expression is TF-specific and we find a striking agreement between the resulting expression curves and the tendency of a TF to appear in homotypic clusters in the human genome, suggesting that intrinsic TFs properties underlie their different representation in homotypic clusters.

## RESULTS

### Accurate measurements of ∼15,000 designed core promoters from a fixed locus in the human genome

We designed a synthetic library of 15,753 oligonucleotides representing native sequences from the human genome including 508 PIC binding regions and 1875 core promoters of coding genes. In addition, we designed synthetic sequences aimed at systematic investigation of the cis-regulatory sequences driving transcription including core promoter elements, 133 TF binding-sites and nucleosome disfavoring sequence. To accurately measure core promoter activity in the genomic context, we developed a high-throughput method for assaying the activity of thousands of sequences from a fixed locus in the human genome using site-specific integration into the “safe harbor” AAVS1 site (**Fig. 1A**)(*17, 18*). Briefly, we obtained a mixed pool of oligonucleotides, 200 basepairs in length, to match our designed sequences and cloned it upstream of an eGFP reporter. We integrated the library into the AAVS1 site in K562 erythroleukemia cells by inducing a double strand break using specific Zinc Finger Nucleases (ZFNs) followed by genomic integration of the reporter cassette by homologous recombination. We then selected cells with a single integrated cassette and used flow cytometry to sort the resulting pool into 16 bins according to eGFP expression normalized by mCherry, which is driven from a constant promoter. In the last step we used deep sequencing to determine the distribution of reads across the different bins for each oligo and extracted the mean and standard deviation to compute mean expression and noise (CV^2^), respectively.

**Figure 1.**
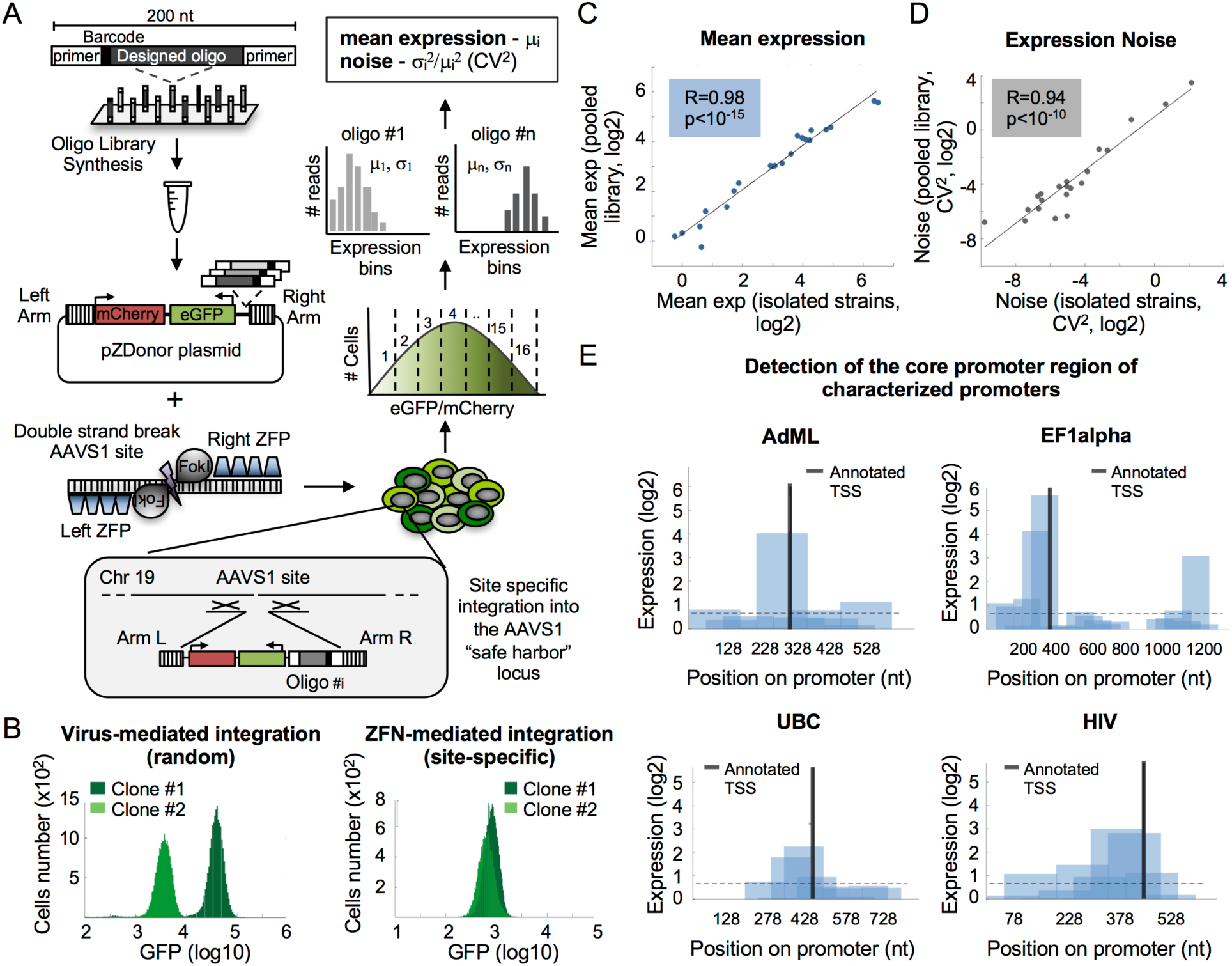
Construction and measurements of 15,753 designed oligonucleotides for core promoter activity using site-specific integration technology. (**A**) 15,753 designed ssDNA oligos in the length of 200nt were synthesized on Agilent programmable arrays and harvested as a single pool. Oligos were amplified by PCR using constant primers and cloned into pZDonor plasmid upstream of eGFP. Plasmids pool was co-nucleofected with mRNAs encoding ZFNs targeting the AAVS1 site into a modified K562 cell line containing only two (of three) copies of the AAVS1 site (see methods). mCherry expression driven from a constant EF1alpha promoter was used to select cells with a single integration by FACS. Cells were then sorted into 16 bins according to eGFP/mCherry ratio. Oligos were amplified from each bin and submitted for deep sequencing. Finally, the distribution among expression bins was determined for each oligo and mean expression and noise were computed. CV - coefficient of variation. (**B**) Comparison between site-specific and random integration. (Left) H1299 cells were infected with retroviruses expressing GFP from a constant promoter. (Right) K562 cells were co-nucleofected with mRNA encoding ZFN-AAVSI and a pZDonor plasmid carrying GFP reporter. Expression levels of two isolated clones in each method are shown. (**C-D**) Accuracy of expression measurements. 21 clones, each expressing a single oligo, were isolated from the library pool and identified by Sanger sequencing. eGFP/mCherry ratio was measured for each clone individually by flow-cytometry. Shown is a comparison between these isolated measurements and those calculated from the pooled expression measurements for mean expression (panel C, R=0.98, Pearson correlation, p<10^-15^) and noise (panel D, R=0.98, Pearson correlation, p<10^-10^). (**E**) Detection of autonomous core promoter activity. Sequences of four fulllength promoters were partitioned *in-silico* into 153nt fragments with large overlap of 103nt between oligos. The characterized TSS is denoted. Dashed line represents the activity threshold determined by the empty vector measurements (methods).

To assess the accuracy of our measurements from site-specific integration in comparison to a traditional retrovirus-based technique, we integrated a single promoter construct multiple times using each system. As expected, in our ZFN system, where all constructs are integrated into the same genomic location, the variability between cells was lower than in the retroviral system, where integration occurs at random locations, spanning a range of ∼1 and ∼2 orders of magnitude in expression, respectively (p<10^-20^ Fig. S1). Moreover, the expression of independently isolated clones was highly similar in the ZFN system, whereas it differed more in the retroviral system (**Fig. 1B**). To evaluate the accuracy of our assay in comparison with each oligo’s individual measurement, we isolated 21 clones from the library pool and measured the expression of each isolated clone using flow cytometry. We found excellent agreement between these measurements and those extracted from the massively parallel assay for both mean expression (R = 0.98, p<10^-15^, **Fig. 1C**) and noise (R=0.94, p<10^-10^, **Fig. 1D**). To gauge the reproducibility of our measurements we designed replicates for different promoters with 10 unique barcodes. For each promoter we examined the distribution of deep sequencing reads among the 16 expression bins for all 10 barcodes, for which synthesis, cloning, sorting and sequencing were independent and found very good agreement between different barcodes (Fig. S2). Finally, to test our ability to measure autonomous core promoter activity we designed 153nt-long sequences tiling the entire length of previously characterized promoters with 103 basepair overlap between oligos. Remarkably, our assay accurately detects the core promoter region in 10 of 11 promoters for which transcription start sites (TSSs) were previously reported. (**Figs. 1E** and S3, Table S1).

Together, these results demonstrate that our method enables highly accurate measurements of core promoter activity for thousands of fully designed sequences in parallel from a fixed location within the human genome.

### Functional measurements of PIC binding sequences in promoters and enhancers

Emerging evidence from recent studies suggests that in contrast to the decades-long wide held belief, transcription initiation is not restricted to promoters. Nascent RNA measurements uncovered thousands of transcription start sites in promoters and enhancers with similar architecture(*19*). Moreover, a genome-wide binding assay identified thousands of PIC-bound regions across the human genome including enhancers(*20*). However, several questions remain unclear, including whether PIC binding sequences can act as functional core promoters, what is the relationship between binding levels and core promoter activity, and whether divergent transcription is a result of true bidirectionality or two adjacent unidirectional initiation sites.

To investigate the functional activity of PIC binding sequences across the human genome, we designed synthetic oligos to match 508 reported binding regions(*20*) and tested their ability to initiate transcription in our reporter assay (**Fig. 2A,** Table S2). Our measurements uncover a positive relationship between PIC binding levels and functional core promoter activity such that regions for which PIC binding is higher also drive stronger expression (p<10^-15^, **Fig. 2B**). To compare the functional transcriptional activity in promoters and enhancers directly we designed oligos to match the sequences bound by PIC from the two regions. To control for potential differences in expression resulting from PIC binding levels, we selected sequences from the same range of binding scores for the two groups. Notably, PIC binding sequences from promoters present higher functional activity than enhancers (p<10^-5^, **Fig. 2C**) that do not stem from differences in binding intensity (p>0.1, **Fig. 2C**).

**Figure 2.**
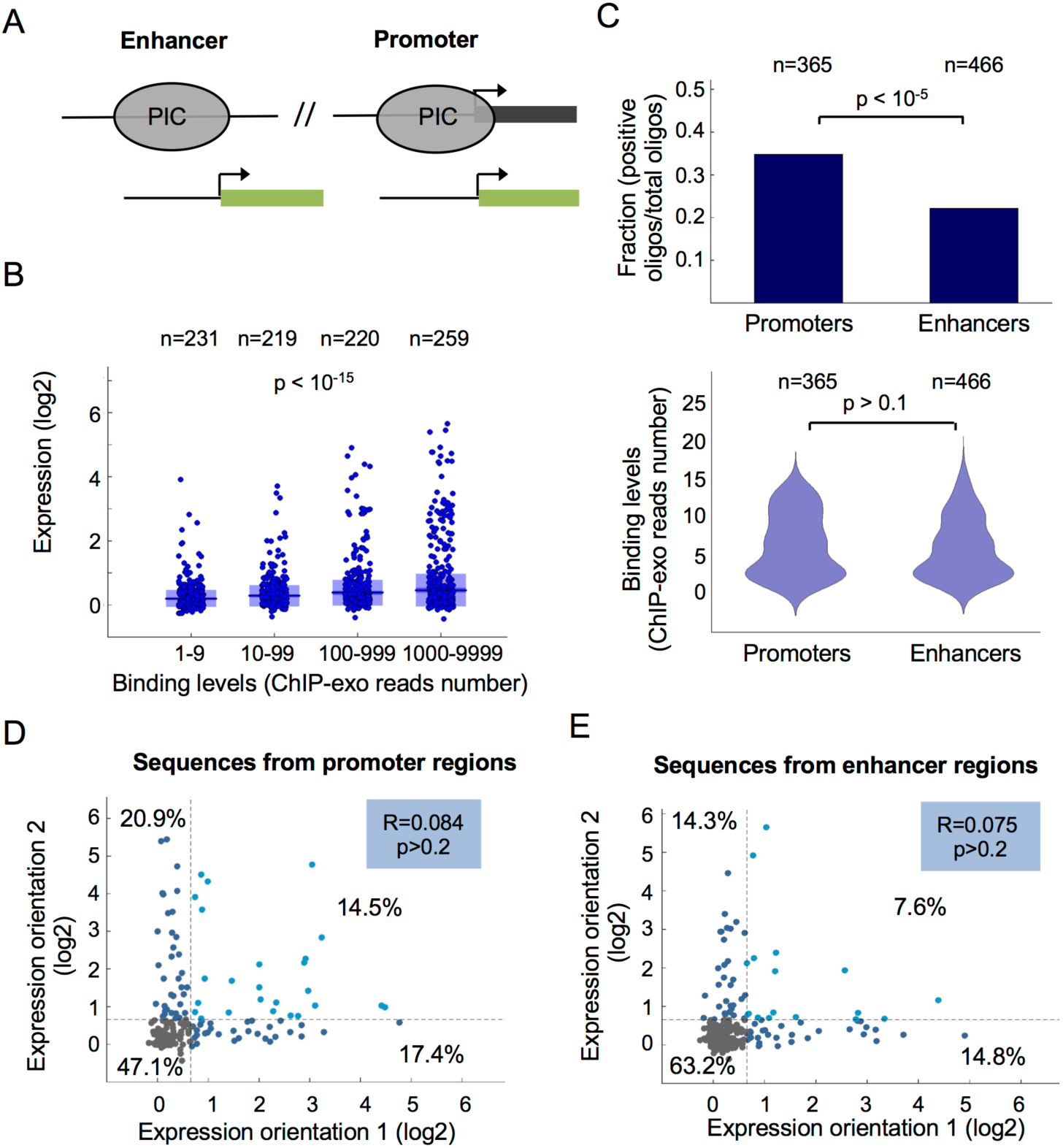
Functional measurements of autonomous core promoter activity of PIC binding sequences from promoters and enhancers. (**A**) Illustration of the designed sequences matching 508 PIC binding regions in promoters and enhancers that were identified by ChIP-exo measurements in K562 cells(*20*). (**B**) Comparison between core promoter activity of sequences with different PIC binding levels. Data was binned into four groups according to the number of ChIP-exo reads and expression measurements were compared between bins (p<10^-15^, Kruskal-Wallis test). (**C**) Comparison between the fraction of positive core promoters for PIC binding sequences from promoters and enhancers (top, p<10^-5^, two-proportion z-test). To avoid biased in activity stemming from different PIC binding levels, sequences with the same number of ChIP-exo reads were selected (bottom, p>0.1, Wilcoxon rank-sum test). (**D-E**) Comparison between core promoter activity of PIC binding sequences from promoters (D) and enhancers (E) in two orientations. Each dot represents a distinct PIC binding sites and expression measurements of designed sequences in two orientations are shown. Dashed lines represent the activity threshold as determined by empty vector measurements and the percentages of cells in each region are denoted. No correlation was detected between expression measurements in the two orientations (R=0.084, p>0.2, for promoters and R= 0.075, p>0.2, for enhancers, Pearson correlation).

Next, we set to investigate whether PIC binding sequences can drive bidirectional transcription. To this end, for each binding site we designed two oligos representing the core promoter sequence (-103 to +50) on either the plus or minus strand. Remarkably, we find no correlation between the expression levels of the two orientations for sequences from promoters (R=0.084, p>0.2, **Fig. 2D**) and enhancers (R=0.075, p>0.2, **Fig. 2E**). Moreover, our measurements uncover that most of the PIC binding sequences display positive activity in only one of the two orientations tested with 38.3% and 29.1% unidirectional vs 14.5% and 7.6% bidirectional expression from promoter and enhancer regions, respectively.

Together, our results demonstrate positive relationships between PIC binding and core promoter activity, an intrinsic difference between sequences from promoters and enhancers, and that core promoters drive unidirectional transcription.

### Systematic investigation of core promoter elements in synthetic and native sequences

The most commonly known core promoter elements are the TATA-box, initiator (Inr), upstream and downstream TFIIB recognition elements (BREu and BREd), motif ten element (MTE), and downstream core promoter element (DPE)(*4*). However, there are no universal core promoter elements that are present in all promoters. Rather, different core promoters exhibit distinct properties that are determined by the presence or absence of particular core promoter motifs.

To systematically test the effect on expression of different core promoter elements we designed synthetic sequences with all possible combinations of the six common elements (**Fig. 3A**)(*3*). We placed the consensus sequences for each of the six elements in five different backgrounds resulting in 320 synthetic oligos (Tables S3-S5). Sorting the tested configurations according to expression, we identify patterns of elements that are abundant in high or low expressing oligos (**Fig. 3A**). To quantitatively assay the separate contribution of each element, we compared the expression levels of all the tested configurations with and without each of the motifs (**Fig. 3B**). Of the six elements tested the only two sequences that led to a significant increase in expression were the TATA-box and the Initiator with 45% and 28% increase, respectively (p<10^-5^ and p<10^-3^, **Fig 3B**). Notably, we found that both the BREu and the BREd elements significantly decreased expression by 35% and 20%, respectively (p<10^-3^ and p<10^-2^, **Fig 3B**). The DPE and MTE elements, which were characterized mostly in *Drosophila*, had no detected effect on expression (p>0.1 and p>0.2. respectively, **Fig 3B**) suggesting that they do not play a substantial role in humans or require additional context-dependent features.

**Figure 3.**
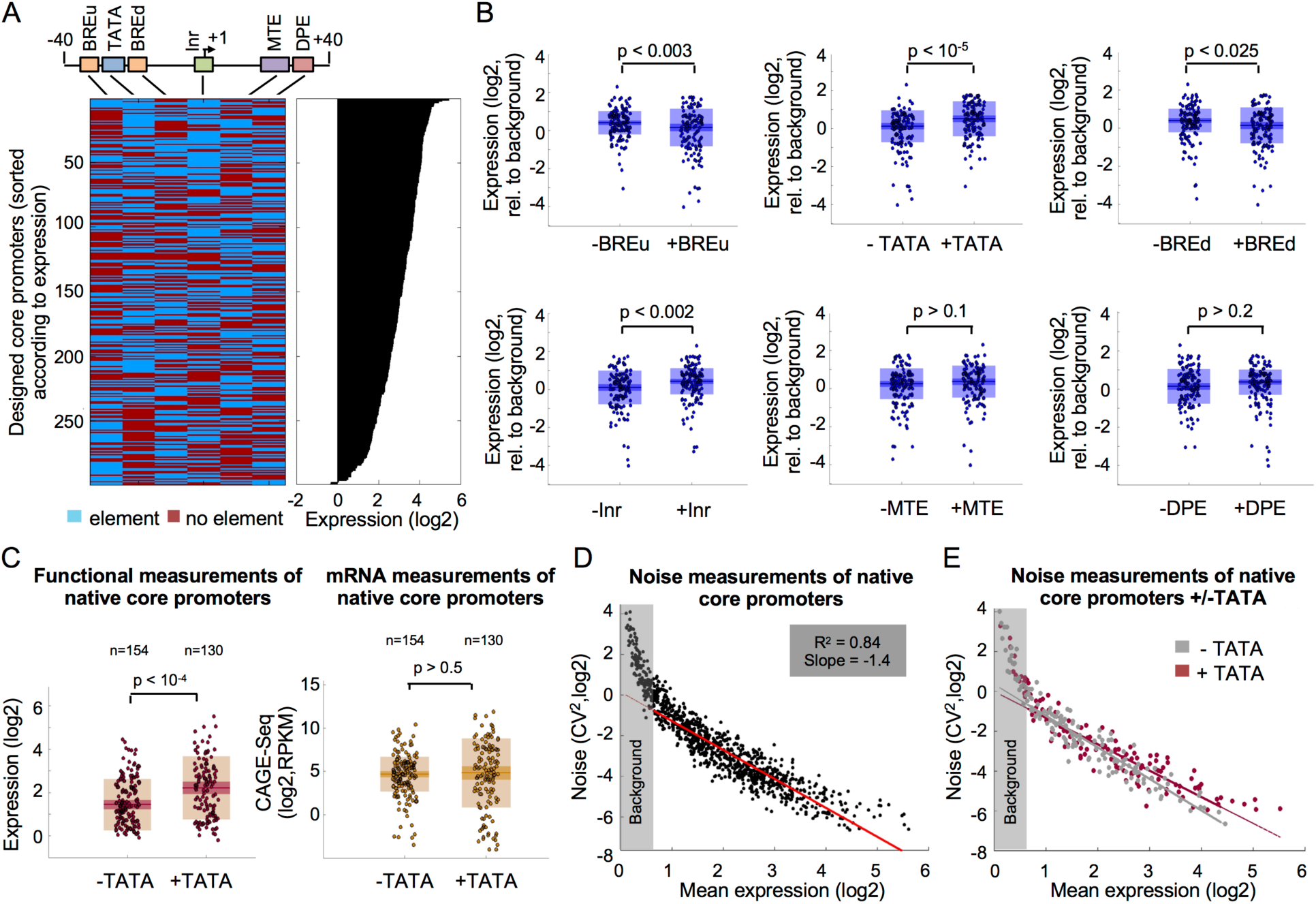
Systematic investigation of six core promoter elements in synthetic configurations and native core promoters from the human genome. (**A**) 320 synthetic oligos representing all possible combination of six core promoter elements on five different backgrounds were designed. Each line in the heatmap (left) represents a single designed oligo and each column represents one of the six elements tested. The configurations were sorted according to the expression measurements (right). (**B**) Comparison between the expression of all the designed sequences with and without each of the six core promoter elements. Wilcoxon rank-sum test was performed to determine significant differences in expression and p-values are denoted. (**C**) The effect of TATA-box in native human core promoters. (left) expression measurements from our functional assay of native core promoters from the human genome with and without a consensus TATA-box. Elevated expression is obtained for promoters with TATA element (p<10^-4^, two-sample t-test). (right) CAGE-seq measurements in K562 cells for the same promoters(*21*). No significant difference was detected between the two groups (p>0.5, two-sample t-test). (**D**) Noise measurement of 990 native core promoters from the human genome as a function of mean expression. A linear fit was performed on oligos with positive core-promoter activity as described before(*24*). (**E**) Comparison of noise measurements of native core promoters with and without a TATA-box.

Although synthetically designed oligos have a tremendous advantage in the investigation of cis-regulatory elements in a controlled setting, their sequences diverge from native promoters in the human genome. To measure the expression of native sequences we designed 1875 core promoters of coding genes using CAGE-seq measurements to determine their TSS(*21*) (Table S6). Next, we set to investigate the effect of the TATA-box in native context using these measurements. Notably, comparing the expression levels of hundreds of native core promoters with and without a consensus TATA-box, we find a significant increase in TATA-containing core promoters (p<10^-4^, **Fig. 3C**). Remarkably, comparing the CAGE-seq measurements, which indicate the transcript levels produced from the native genomic locus, we find no significant difference between the two groups (p>0.5, **Fig. 3C**). This finding demonstrates the importance of performing designated functional assays to decipher the autonomous activity of core promoters when isolated from additional factors influencing the transcriptional output such as neighboring enhancers and local chromatin environment.

In addition to regulating mean expression, core promoter elements such as the TATA-box were also shown to have an effect on cell-to-cell variability in yeast, or expression noise(*22, 23*). To investigate the effect of the core promoter sequence on noise we used the distributions of reads across the expression bins to compute for each oligo the mean and standard deviation. We quantified the noise by the squared coefficient of variation (CV^2^), that is the variance divided by the square mean(*24*) (**Fig. 1A**). Notably, we find that noise is scaled with mean expression with similar dependency as described for yeast(*24*) (fitted slope of −1.4, **Fig 3D**). However, in contrast to yeast promoters(*25*) we do not find large differences in noise for the same mean expression with most of the variability explained by the mean expression (R^2^=0.84, **Fig. 3D**). Moreover, we do not find substantial differences between TATA and TATA-less sequences in native core promoters (**Fig. 3E**) or for any of the six core promoter elements tested in the synthetic sequences (Fig. S4). A potential source for the observed difference between yeast and human cells is the generation time. While yeast cells divide every ∼1.5 hours, the generation time of most cultured mammalian cells is ∼24 hours. Thus, using stable eGFP reporter, as done in our assay, can buffer the effect of rapid fluctuations in mRNA levels(*26*). However, since the median half-life of mammalian proteins is 46 hours(*27*), the stable eGFP reporter that we use here may better represent the true cell-to-cell variability of most of the endogenous proteins.

Taken together, our findings demonstrate significant effects of core promoter elements on mean expression, with positive effects for the TATA and the Inr, and negative effects for the BRE upstream and downstream elements.

### TATA and Inr additively increase expression at preferable distances

A key question in the investigation of core promoter elements is the effect of motif combinations on expression. Bioinformatic analyses suggested that core promoter elements act in a synergistic manner to recruit RNA Pol II(*28*). Moreover, previous studies demonstrated that their coordinated effect on transcription depends on the distance between the elements(*29, 30*).

To investigate the relationship between the TATA and the Inr, both found to positively regulate promoter activity in our assay, we compared the expression of all tested configurations with TATA to those containing TATA and Inr. We found that adding Inr to TATA-containing promoters results in increased expression (p<10^-3^, **Fig. 4A**). Next, we set to investigate if the two elements act in synergy by comparing the expression levels of oligos containing both elements to the sum of expression of oligos containing TATA or Inr separately (**Fig. 4B**). Interestingly, we do not find higher expression for oligos with the two elements suggesting that they act in a partially additive manner and not synergistically to increase transcription.

**Figure 4.**
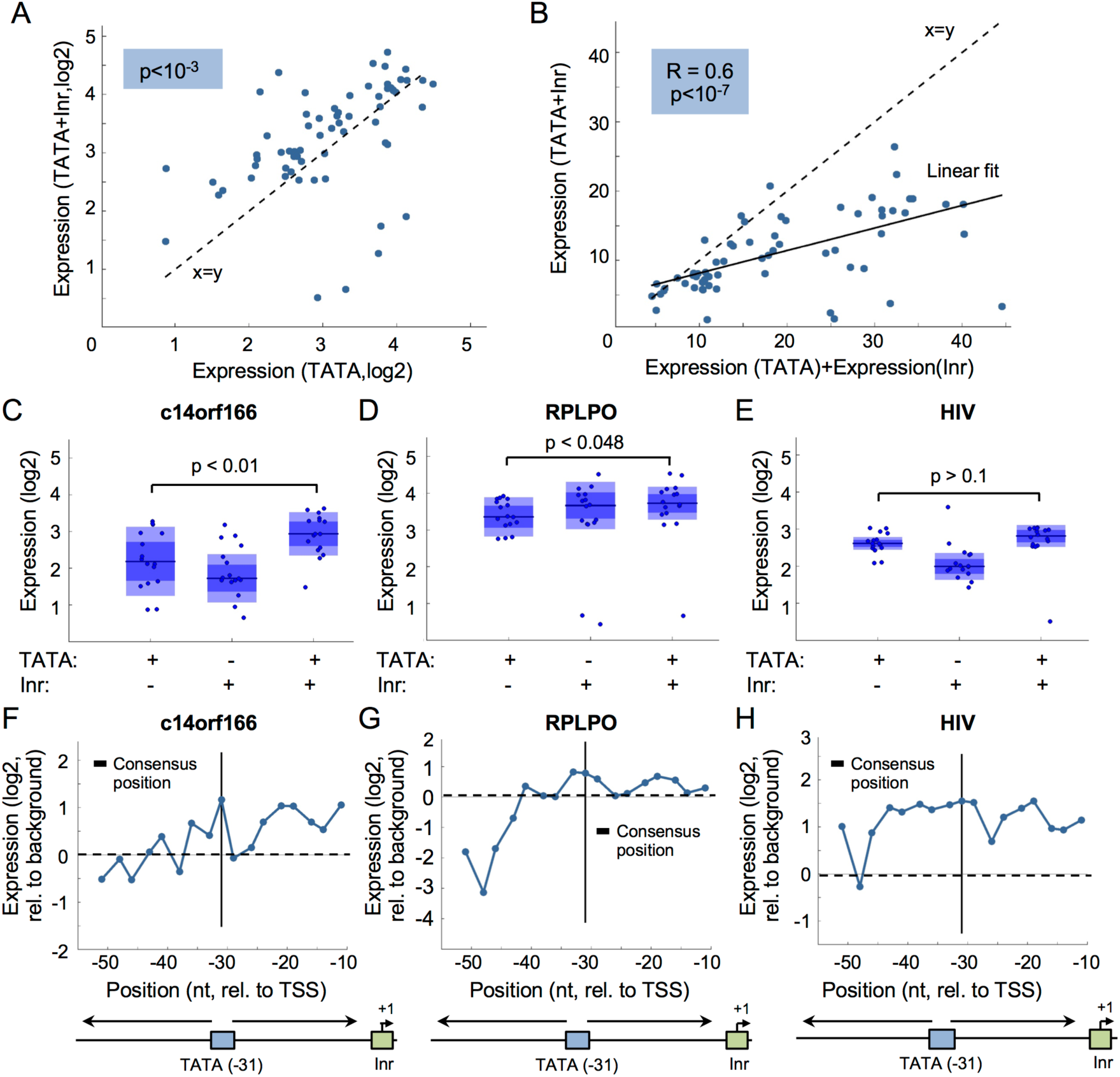
The effect on expression of TATA-box and Initiator combinations and relative distance in different backgrounds. (**A**) Comparison of expression levels of synthetic oligos with TATA to those containing both TATA and Inr. Each dot represents a pair of sequences with either TATA or TATA+Inr elements. An increase in expression is observed when adding Inr (p<10^-3^, Wilcoxon signed rank test). (**B**) Testing for synergy between the TATA and the Inr elements. Each dot represents a pair of expression values. On the x-axis, expression was computed as the sum of the expression of separate oligos with either TATA or Inr. The y-axis represents expression measurements of oligos that contain the two elements. (**C-E**) Comparison of oligos with either TATA, Inr and TATA+Inr in three different promoter backgrounds. Presented p-values were computed by Wilcoxon rank-sum test. (**F-G**) Testing the effect of the distance between the TATA and the Inr in three different backgrounds. We designed oligos in which we placed the Inr in its consensus position and systematically changed the location of the TATA (2-3nt increment). Each blue dot represents the expression levels at a single position. The consensus position of the TATA (-31) is denoted.

To test whether the activity of core promoter elements depends on the background sequence, we analyzed the effects on expression of the TATA and the Inr in three different backgrounds separately. Notably, our results show that while for some backgrounds (C14orf166 and RPLPO) expression increases when adding Inr to TATA-containing oligos (p<0.01 and p<0.05, **Figs. 4C,D**) for others (HIV) adding an Inr does not increase expression beyond the effect of the TATA (p>0.1, **Fig. 4E**). Remarkably, promoters for which adding Inr to the TATA leads to an increase in expression also present greater sensitivity to the distance between the two elements in general with maximal expression achieved when placed in the consensus reported position (-31) (**Figs. 4F,G,** Table S7). In contrast, the HIV promoter, for which adding Inr to the TATA had no significant effect, was more robust to changes in the TATA location with similar expression levels for the majority of the positions tested (**Fig. 4H**).

Notably, in all three backgrounds, expression is higher when the TATA is placed around positions −10, −20 and −30 relative to the TSS than positions −15 and −25 (**Figs. 4F-H**). This ∼10bp periodicity, which matches the DNA double helix geometry, implies that the stereospecific alignment between the TATA and the TSS is important for expression, as was previously described for transcription factors(*2, 10, 31, 32*). Interestingly, periodicity was not observed in the CMV background (Fig. S5) suggesting that the sequence in which the elements are embedded affects alignment-dependent interactions.

Together, our results show that the TATA and the Inr elements can act additively to enhance transcription at preferable distances that facilitate stereospecific alignment between the two elements.

### Comprehensive activity screen for 133 TF binding-sites and nucleosome disfavoring sequences

In addition to the core promoter elements, the recruitment of the pre-initiation complex is regulated by specific TFs that bind the proximal promoter region. Computational and high-throughput experimental approaches had characterized binding specificity(*33*) and mapped the positions of TF binding-sites in the human genome(*34–36*). However, since the expression levels of TFs, their localization and post-translational modification vary between cell types, we cannot determine which TF binding-sites will affect expression and to what extent.

To directly survey the activity levels of TFs, we designed promoters in which we planted four copies of each of 133 binding-sites for 70 different TFs in two different backgrounds (**Fig. 5A**, Table S8). To test the effect of directionality we placed the sites in either the forward or reverse orientation. We found positive activity for 63% and 58% binding-sites in the Beta-Actin and the CMV backgrounds, respectively, spanning a dynamic range of ∼30-fold in expression (**Fig. 5B**). Notably, expression levels in both orientations are highly correlated (R=0.81, p<10^-20^, **Fig. 5C**) suggesting that TF-driven expression is not sensitive to the binding-site directionality. Similarly, we find good agreement between expression measurements in the two tested backgrounds (R=0.72, p<10^-20^, **Fig. 5D**).

**Figure 5.**
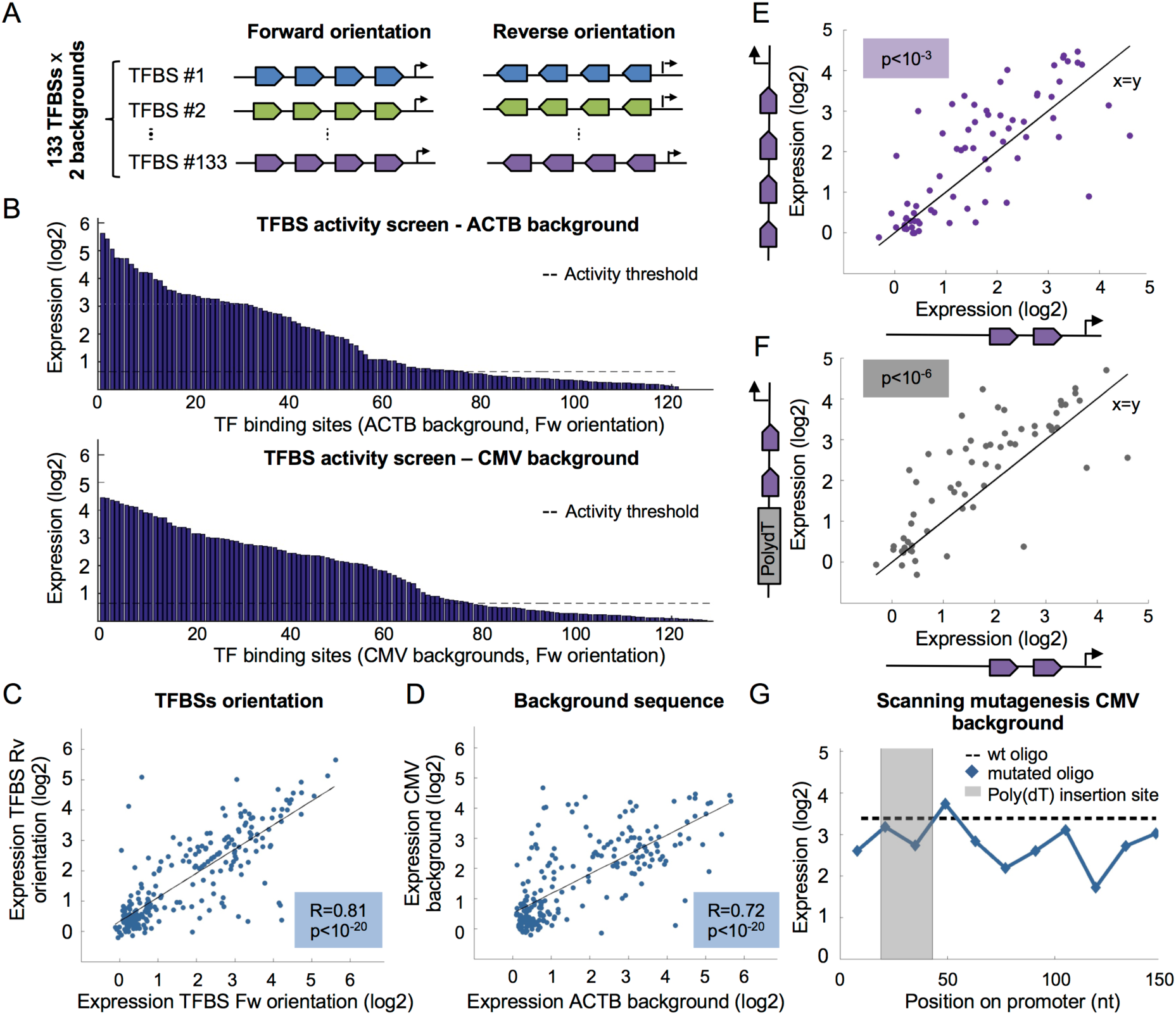
TF activity screen for 133 binding-sites and the effect of nucleosome disfavoring sequence on expression. (**A**) Illustration of designed oligos for TF activity screen. 133 binding-sites for 70 TFs were placed in four copies in either the forward or the reverse orientation in two backgrounds. (**B**) Expression measurements of oligos containing forward TF binding-sites in two different backgrounds. Each bar represents a single binding-site. Activity threshold determined by the empty vector is denoted. (**C**) Comparison between expression measurements of binding-sites in two orientations. Each dot represents a pair of sequences for the same binding site when places in the forward or the reverse orientation (R=0.81, p<10^-20^, Pearson correlation). (**D**) Comparison between expression measurements of binding-sites in different backgrounds. Each dot represents a pair of sequences for the same binding site when places in the Beta-Actin or the CMV backgrounds (R=0.72, p<10^-20^, Pearson correlation). (**E**) Testing the effect on expression of adding two TF binding-sites. Each dot represents a pair of designed promoters with either two or four sites for one of the 70 TFs tested in the CMV background. An increase in expression is observed for most TFs (p<10^-3^, Wilcoxon signed rank test). (**F**) Testing the effect on expression of nucleosome disfavoring sequence. 25-mer poly(dA:dT) tract was added upstream to two binding-site for 70 TFs. An increase in expression is observed for most TFs (p<10^-6^, Wilcoxon signed rank test). (**G**) Systematic scanning mutagenesis to identify cis-regulatory elements in the CMV promoter. 11 mutated oligos were designed, each contains a 14nt window in which all nucleotides were mutated. Each dot represents expression of one mutated oligo. No elevation in expression is observed when mutating the sequences in which the poly(dA:dT) was inserted.

Previous studies from our lab demonstrated that transcription in yeast can be elevated either by increasing the number of TF binding-sites or by adding poly(dA:dT) tracts that act as nucleosomes repelling sequences both in vivo and in vitro (*10, 37–40*). To investigate these effects in human for a large number of factors, we designed promoters with two binding-sites for 70 TFs in two backgrounds. We then placed either two additional binding-sites or poly(dA:dT) tracts 25 basepairs in length upstream to the two existing sites. As expected, we find increase in expression when adding two TF binding-sites to the CMV background (p<10^-3^, **Fig. 5E**). Notably, poly(dA:dT) tracts led to increase in expression for most of the TFs tested, similar to what we reported for yeast promoters(*37, 41*) (p<10^-6^, **Fig. 5F**). To ensure that the obtained increase in expression is not a result of destruction of a repressive sequence in the promoter background, we mapped the cis-regulatory elements using systematic mutagenesis and found no increase in expression when introducing random mutations in the same region (**Fig. 5G**). Testing the Beta-Actin promoter we did not find a general increase in response to poly(dA:dT) tracts (Fig. S6). However, the same background was also not affected by additional TF binding-sites suggesting that for some TFs the expression driven by two sites is nearly saturated so that the contribution of additional elements cannot be accurately evaluated.

Together, our results, constituting the largest profiling of TF activity in human cells to date, demonstrate bidirectional activity and show that similar to what was shown in yeast, poly(dA:dT) tracts can sometimes increase expression in similar levels to TF binding-sites.

### The effect of binding site number on expression is TF-specific

Proximal promoters and distal enhancers are enriched for multiple sites for the same factor, also known as homotypic clusters of TF binding-sites (HCT). Their conservation in vertebrates and invertebrates suggests that this is a general organization principle of cisregulatory sequences(*42*). Studies that examined the number of homotypic clusters for different TFs in the human genomes found a wide range of behaviors, with some factors (e.g., SP1) forming a large number of HCTs while others (e.g., CREB) are rarely found in homotypic clusters(*42*). This observation suggests that the effect on expression of multiple sites for the same factor depends on the identity of the TF, resulting in different relationships between binding site number and expression (**Fig. 6A**).

**Figure 6.**
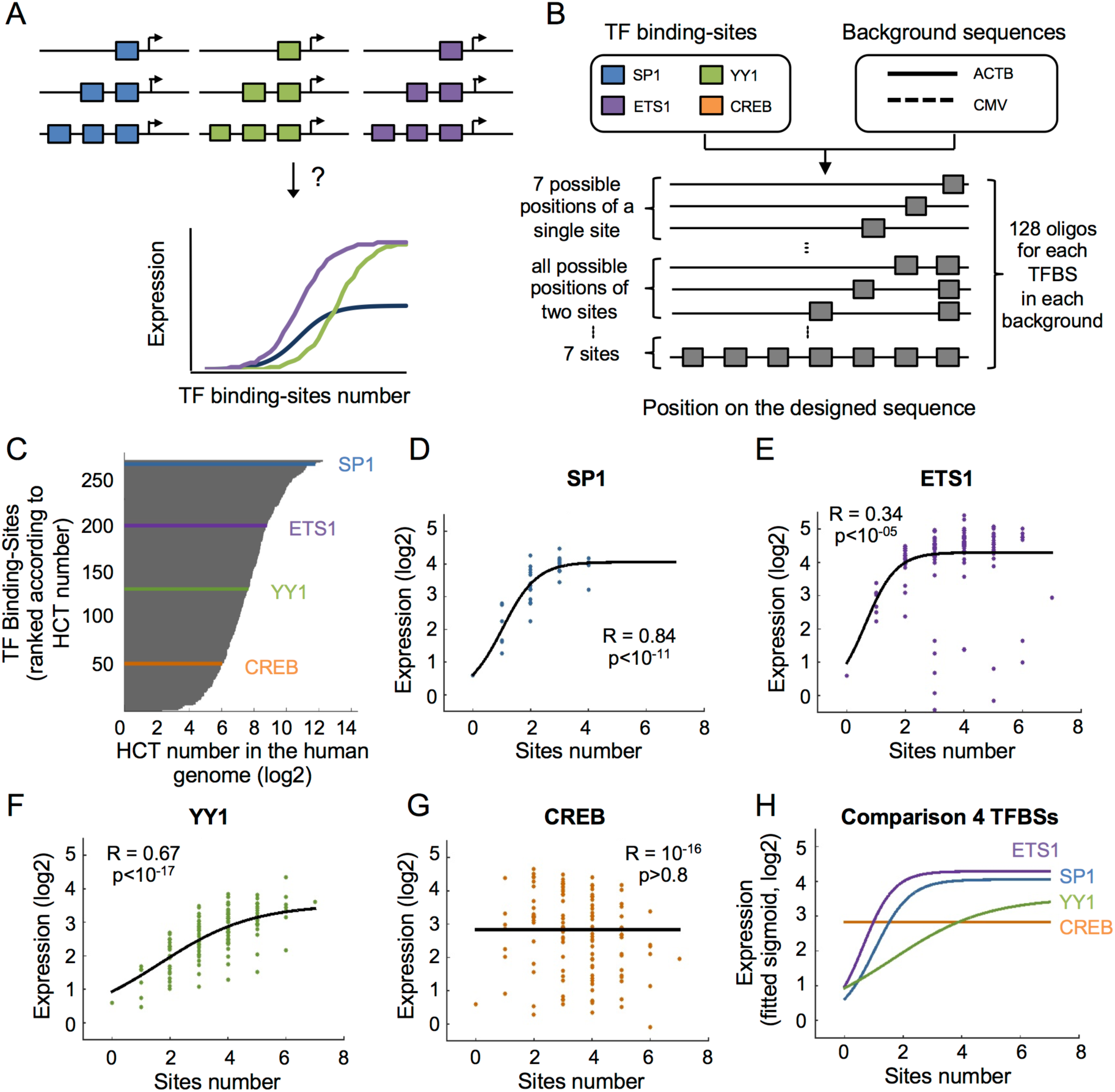
Systematic interrogation of the effect of homotypic TF binding-sites number on expression. (**A**) Illustration of different expression functions when adding homotypic binding sites for different TFs. (**B**) The design of 1,024 synthetic oligos to systematically investigate the effect of sites number on expression. Four different TFs were planted in all possible combinations of 1-7 sites in 7 predefined positions within two different background sequences. (**C**) Shown is the number of homotypic clusters for TF bindingsites (HCTs) of different factors in the human genomes. Data was analyzed from Gotea *et al*.(*42*). Denoted are the four TFs chosen for the design of the synthetic oligos representing different numbers of HCT. (**D-G**) Expression measurements of oligos with increasing number of sites for SP1 (D), ETS1 (E), YY1 (F) and CREB (G) in the Beta-Actin background. Each dot represents a single oligo in the library. A logistic function was fitted (methods) and the correlation between the expression measurement and the fitted values are shown for each TF. (**H**) A summary plot of the four expression curves that were computed in D-G for direct comparison between TFs.

To systematically interrogate the effect of homotypic site number on expression we designed oligos in which we separately planted the sequences of four different TF binding-sites in 1-7 copies. To control for the effects of the binding site distance from the TSS, the distance between adjacent sites and the immediate flanking sequence, we planted each TF binding site in all possible combinations of 1-7 sites at 7 predefined positions. We tested two backgrounds resulting in a total of 1,024 oligos (**Fig. 6B,** Table S9). We selected four factors that are common in the proximal promoter region with different number of endogenous homotypic clusters in the human genome (**Fig. 6C**). To evaluate the relationship between binding site number and expression, we fitted a logistic function to the expression measurements (**Figs. 6D-G**, methods). Comparing the four TFs in the Beta-Actin background, we find a striking agreement between the number of homotypic sites in the human genome and the obtained expression curves (**Figs. 6C,H**). Specifically, SP1 and ETS1, which have the highest number of homotypic sites of the four factors tested (3522 and 448, respectively), present the steepest increase (slopes of 1.67 and 1.91, respectively) and achieve the highest maximal expression levels (4.06 and 4.29, respectively) (p<10^-11^ and p<10^-05^, **Figs. 6D,E,H**). YY1, which has an intermediate number of homotypic sites (202), presents moderate increase (slope=0.62) and intermediate maximal expression levels (3.53) (p<10^-17^, **Figs. 6F,H**). Finally, increasing the number of sites for CREB, which has the lowest number of homotypic sites (66), had no significant effect on expression (R=0, p>0.8, **Fig. 6G,H**). Testing the expression curves in the CMV background we find similar trend with three of the four factors preserving the same rank as in the Beta-Actin background (Fig. S7). Remarkably, here too we found that adding CREB sites does not increase expression (R=0.17, p>0.05).

Taken together, our findings demonstrate that the effect of homotypic sites on expression is factor-specific and that TFs that are naturally more prevalent in homotypic clusters in the genome also display higher dependency between binding-site number and expression.

## DISCUSSION

Here we established a high throughput experimental system to investigate thousands of designed sequences in a controlled genomic setting. We used this method to perform a comprehensive and systematic study of the core promoter and the proximal promoter regions in human. Our measurements reveal the functional activity and directionality of genomic DNA sequences bound directly by the pre-initiation complex in promoters and enhancers. Investigating native and synthetic sequences, we characterize the cis-regulatory elements underlying transcription including core promoter elements, TF binding-sites and nucleosome disfavoring sequence. By examining a large space of configurations and distances we show how these elements combine to orchestrate a transcriptional output. We provide activity measurements of 133 TF-binding sites and quantify the effect on expression of multiple homotypic sites for different TFs. Importantly, we show that interaction between elements, either core promoter elements or TF binding-sites, are not universal and can vary between different backgrounds or factors. In turn, these differences can be translated into organizational principles of regulatory regions in the human genomes.

A growing number of studies employ massively parallel reporter assays (MPRAs) to decipher gene expression regulation at the levels of transcription, translation and mRNA stability(*11, 15, 43–46*). These types of experiments emphasize the need for accurate methodologies aiming at investigating designed sequences from a native genomic context. In this study, we developed a method for measuring thousands of designed oligos from a fixed location within the human genome with high efficiency and accuracy. Our method can readily be adapted to assay different types of regulatory elements providing a valuable tool to interrogate gene expression. Importantly, site-specific integration followed by flowcytometry sorting provides single cell information allowing for systematic investigation of cell-to-cell variability that cannot be inferred from current MPRA methods, in which each cell is transfected with multiple different constructs. Thus, our method enables multidimensional investigation of the effect of sequence on expression at the populationand single cell-levels, allowing to infer both mean expression and noise in a single experiment.

Our findings shed additional light on the divergent nature of human promoters. The discovery of bidirectional transcription by genome-wide measurements of nested transcripts led to ongoing discussion on the existence and mechanisms underlying bidirectional promoters(*47, 48*). In a recent study, Core *et al.* investigated the landscape and architecture of TSSs across the human genome and found that divergent transcription in both promoters and enhancers is facilitated by two distinct core promoters separated by ∼110bp(*19, 49*). Interestingly, functional measurements of random genomic fragments using massively parallel reporter assay identified divergent promoter activity(*5*). However, since the assayed sequences were in the length of 0.2-2kb, one cannot tell whether the activity observed represents “true” divergent sequences or two adjacent unidirectional core promoters. To address this question directly, we designed oligos to specifically match the core promoter region by taking 103 basepairs upstream and 50 basepairs downstream of hundreds of PIC binding sites. Notably, our functional measurements uncover that core promoters mostly drive unidirectional transcription. Moreover, in the model proposed by Core *et al*., a centered TF directs the pre-initiation complex to initiate transcription from the two core promoters. Remarkably, in line with this model, we find high agreement between the activity levels of 133 TF binding-sites when placed in the forward and the reverse orientations. Together, our study provides direct functional measurements supporting a model by which divergent promoter activity is driven by two distinct unidirectional core promoters sharing bidirectional TF binding-sites.

Our systematic investigation of all possible combinations of the common six core promoter elements in various backgrounds reveals that while the TATA and the Inr increase expression, the BRE upstream and downstream elements lead to reduction in core promoter activity. Interestingly, BREu and BREd have been found to elicit both positive and negative effects on basal and TF-induces transcription(*4, 50–53*). Considering that both BREs can act to stabilize the assembly of the pre-initiation complex through interactions with the TFIIB general transcription factor (GTF), their negative effect on expression may be counter intuitive. However, it was suggested that while GTF–core promoter interactions can enhance the formation of the pre-initiation complex, they might also impede the transition from initiation to promoter escape(*50*). Thus, sequence elements that increase the affinity between the initiation complex and the core promoter can have a negative impact on the transcriptional outcome.

TF binding-sites can appear in homotypic and heterotypic clusters in the genome. An intriguing question is which of these organizational principles results in higher expression. Interestingly, a recent study that assayed 12 liver-specific transcription factors in homotypic and heterotypic clusters found that heterotypic elements are in general stronger than homotypic ones(*54*). However, since TFs differ in their DNA-binding, transactivation and oligomerization domains, they may not adhere to one universal rule. Indeed, examining the human genome reveals that the tendency to appear in homotypic clusters is not uniform across TF binding-sites(*42*). Our systematic measurements of multiple homotypic sites for distinct TFs uncover differences between their expression curves. Thus, TFs may employ different strategies to enhance transcription and while some can “benefit” from homotypic sites for others combining with heterotypic site may result in higher expression. In addition, we find a striking agreement between the TF-specific expression curves resulting from multiple homotypic sites and the corresponding representation of a TF in homotypic clusters across the human genome. This finding suggests that intrinsic differences between TFs may be encoded in the genome and that we can use this information to increase our understanding of the various activation patterns of TFs.

Genomic analysis of tumours using next generation sequencing (NGS) has led to the identification of thousands of mutations, many of which reside within non-coding sequences. However, the effect of DNA sequence changes in regulatory regions remains elusive. Importantly, recent studies of breast cancer and melanoma had shown that the acquisition of mutations in the promoter sequence of four genes alter their expression by affecting TF binding-sites and proteins recruitment to the promoter region(*55, 56*). Thus, deciphering the mapping between promoters’ architecture and gene expression is key for understanding the transcriptional events underlying the development of cancer and additional genetic diseases. Our comprehensive characterization of human promoters including the directionality of core promoters and TF binding-sites; the portrayal of core promoter elements that positively and negatively regulate transcription; and the effect of elements combinations and distances, adds new insights into rules of transcriptional regulation. In turn, these insights can facilitate the interpretation of the hundreds of DNA sequence changes associated with multiple diseases.

## MATERIALS AND METHODS

### 1. Experimental Procedures

#### 1.1 Cell culture

K562 cells (CCL-243, ATCC) were cultured in tissue culture flasks (Nunc) in Iscove’s medium (Biological Industries, Beit-Haemek, Israel (BI)) supplemented with 10% fetal bovine serum (BI) and 1% penicillin and streptomycin (BI). H1299 human lung carcinoma cells with ecotropic receptor were cultured in RPMI 1640 medium (Gibco), supplemented with 10% fetal bovine serum (BI), and 1% penicillin and streptomycin (BI). Phoenix virus packaging cells were cultured in DMEM medium, supplemented with 2 mM L-glutamine, 10% fetal bovine serum (BI), and 1% penicillin and streptomycin (BI). Cells were kept at 37°C in a humidified atmosphere containing 5% CO_2_ and were frozen in complete media with 5-7% DMSO (Sigma).

#### 1.2 Plasmids

pZDonor AAVS1 was purchased from Sigma, as a part of the CompoZr Targeted Integration Kit – AAVS1, as were pZFN1 and pZFN2. pZDonor HindIII was a kind gift from Fyodor Urnov (Sangamo BioSciences). pPRIGp mChHA retroviral vector(*57*) was a kind gift from Patrick Martin (Université de Nice).

#### 1.3 Synthetic library production and amplification

The production and amplification steps were carried out essentially as in (*10*). Agilent Oligo Library Synthesis (OLS) technology was used to produce a pool of 55,000 different fully designed single-stranded 200-mers, a subset of 15,753 of which comprised the synthetic library presented in this study. Each designed oligo contains subset specific priming sites, leaving 164nt for the variable region. The library was synthesized using Agilent’s on-array synthesis technology (*58, 59*) and then provided to us as an oligo pool in a single tube (10pmol). The pool of oligos was dissolved in 200μl TE. 5.5ng of the library (1:50 dilution) were divided into 16 tubes, and each tube was amplified using PCR. The primers used for amplification of the library included sites for the restriction enzymes AscI and RsrII, for cloning into the library master plasmid. The oligonucleotides were amplified using constant primers in the length of 51nt, which are complementary to the subset primer (underlined) and adds the restriction sites (bold) and a tail of approx. 30nt to allow identification of products that were not properly cut by restriction enzymes in the next step. Primers sequences: upstream primer – 5’ – TTGTTCCGCCGCTTCGCTGACTGTG**GGCGCGCC**<u>CGCGTCGCCGTGAGG AGG</u> −3’, downstream primer 5’ – TCAGTCGCCGCTGCCAGATCGCGGT**CGGTCCG**A<u>GCCCCACGGAGGTGC CAC</u> – 3’. Each PCR reaction contained 24μl of water with 0.323ng DNA, 10μl of 5X Herculase II reaction buffer, 5μl 2.5mM dNTPs each, 2.5μl 20uM Fw primer, 2.5μl 20uM Rv primer, and 1μl Herculase II Fusion DNA Polymerase (Agilent Technologies, Santa Clara, California). The parameters for PCR were 95°C for 1 min, 14 cycles of 95°C for 20 sec and 68°C for 1 min, each, and finally one cycle of 68°C for 4 min. The PCR products from all 16 tubes were joined and concentrated using Amicon Ultra, 0.5ml 30K centrifugal filters (Merck Millipore) for DNA Purification and Concentration. The concentrated DNA was then purified using a PCR minielute purification kit (QIAGEN) according to the manufacturer’s instructions.

#### 1.4 Construction of reporter master plasmids

A dual fluorophore master plasmid was constructed, to allow cloning of the library as a proximal promoter of a single fluorophore, while using another fluorophore for normalization. In order to minimize trans-activation between the eGFP driving Beat-Actin (ACTB) promoter, into which the library was cloned, and the EF1alpha promoter driving the mCherry control fluorophore, the master plasmid was designed to maximize the distance between the promoters. Thus, a sequence encoding two cassettes (each containing a promoter, fluorophore and a terminator) placed back to back (with adjacent terminators) was synthesized by Biomatik (Canada) and cloned into the pZDonor plasmid. The eGFP cassette included a fragment of (-468,-122) of the human ActB promoter (from genomic sequence NG_007992.1), followed by sites for AscI and RsrII restriction enzymes, a 5’UTR and the chimeric intron from the pci-neo plasmid (Promega), eGFP gene and the SV40 polyA. A linker sequence of 25bp was designed between the AscI and RsrII restriction enzyme sites (GGGTGTGTTGTTGGTGGGTTGGGTG), and was present instead of the library in the master plasmid control. The mCherry cassette included the EF1alpha promoter, mCherry and the BGH polyA.

#### 1.5 Synthetic library cloning into the master plasmid

The amplified synthetic library was cloned into the master plasmid described above. Library cloning into the master plasmid was adopted from a protocol that was previously described for a lenti-virus based library(*15*). Purified library DNA (720ng total) was cut with the unique restriction enzymes AscI and RsrII (Fermentas FastDigest) for 2 hours at 37°C in four 40 μl reactions containing 4μl FD buffer, 1μl of AscI enzyme, 2.5μl of RsrII enzyme, 0.8μl DTT, and 18μl DNA, followed by a heat inactivation step of 20 min at 65°C. Digested DNA was separated from smaller fragments and uncut PCR products by electrophoresis on a 2.5% agarose gel stained with GelStar (Cambrex Bio Science Rockland). Fragments in the size of 200bp were cut from the gel and eluted using electroelution Midi GeBAflex tubes (GeBA, Kfar Hanagid, Israel)). Eluted DNA was precipitated using standard NaAcetate\Isoprpoanol protocol. The master plasmid was cut with AscI and RsrII (Fermentas FastDigest) for 2.5 hours at 37°C in a reaction mixture containing 9μl FD buffer, 3μl of each enzyme, 3μl Alkaline Phosphatase (Fermentas), and 4.5μg of the plasmid in a total volume of 90μl, followed by a heat inactivation step of 20 min at 65°C. Digested DNA was purified using a PCR purification kit (QIAGEN). The digested plasmids and DNA library were ligated for 0.5 hr at room temperature in two 10μl reactions, each containing 150ng plasmid and the library in a molar ratio of 1:1, 1μl CloneDirect 10X Ligation Buffer, and 1μl CloneSmart DNA Ligase (Lucigen Corporation) followed by a heat inactivation step of 15 min at 70°C. 14μl ligated DNA was transformed into a tube of E.cloni 10G electrocompetent cells (Lucigen) divided to 7 aliquots (25μl each) which were then plated on 28 LB agar (200mg/ml amp) 15cm plates. To ensure that the ligation products only contain a single insert we performed colony PCR on 93 random colonies. The volume of each PCR reaction was 30μl; each reaction contained a random colony picked from a LB plate, 3μl of 10X DreamTaq buffer, 3μl 2mM dNTPs mix, 1.2μl 10μM 5’ primer, 1.2μl 10μM 3’ primer, 0.3μl DreamTaq Polymerase (Thermo scientific). The parameters for PCR were 95°C for 5 min, 30 cycles of 95°C for 30s, 68°C for 30s, and 72°C for 40s, each, and finally one cycle of 72°C for 5 min. The primers used for colony PCR were taken from the ActB promoter (5’ – CTCTTCCTCAATCTCGCTCTCGCTC – 3’) and the chimeric intron (5’ – GACCAATAGGTGCCTATCAGAAACGC – 3’). Out of the 93 colonies evaluated, only 3 had multiple inserts. To ensure that all ∼15,000 oligos are represented we collected over 2·10^6^ colonies sixteen hours after transformation, by scraping the plates into LB medium. Library pooled plasmids were purified using a NucleoBond Xtra maxi kit (Macherey Nagel). Following the purification, the library plasmids were extracted from a 0.8% agarose gel, in order to clean them from free library DNA that presented a toxic effect on library nucleofected cells.

#### 1.6 *in-vitro* transcription of ZFN mRNA

ZFN mRNA was *in-vitro* transcribed from pZFN1 and pZFN2 plasmids (Sigma) according to the manufacturer’s protocol, using MessageMAX T7 ARCA-Capped Message Transcription Kit and Poly(A) Polymerase Tailing Kit (CellScript). The RNA was then purified using MEGAClear kit (Ambion), the concentration was measured and integrity and polyadenylation were verified by high-sensitivity RNA Tapestation (Agilent). Small aliquots (5-10μl) containing 600ng/μl of each of the two ZFN mRNAs were stored in −80°C.

#### 1.7 Preparation of a dual copy AAVS1 site K562 cell line

In order to reduce the number of possible AAVS1 integration sites from the three sites present in K562 cells, cells were nucleofected with ZFN mRNA and a pZDonor plasmid containing a HindIII site between the homology arms. Single cells were sorted by FACS, and grown for up to a month to establish isogenic populations. Cells from the resulting populations were renucleofected with a fluorescent reporter to assess the number of possible genomic integrations. Cell lines exhibiting lower expression of the reporter were selected. In this manner, a cell line in which only two AAVS1 copies were present was retrieved, and was used for all subsequent experiments.

#### 1.8 Nucleofection of library into K562 cells and site-specific integration into the AAVSI locus

The purified plasmid library was nucleofected into K562 cells and genomically integrated using the Zinc Finger Nuclease (ZFN) system for site-specific integration, with the CompoZr^®^ Targeted Integration Kit - AAVS1 kit (Sigma). To ensure adequate library representation, 15 nucleofections with the purified plasmid library were carried out, each to 4 million cells. This number of cells was calculated to result in a thousand transfected cells per each sequence variant and at least 40 single integration events in average per variant. A master plasmid with no insert was also genomically integrated in the same manner. Nucleofections were performed using Amaxa^®^ Cell Line Nucleofector^®^ Kit V (LONZA), program T-16. Cells were centrifuged and washed twice with 20ml of Hank’s Balanced Salt Solution (HBSS, SIGMA), followed by resuspension in 100μl room temperature solution V (Amaxa^®^ Cell Line Nucleofector^®^ Kit V). Next, the cells were mixed with 2.75μg of donor plasmid and 0.6μg each *in-vitro* transcribed ZFN mRNA just prior to nucleofection. A purified plasmid library was also nucleofected without the addition of ZFN to assess the background level of non-specific integration and the time for plasmid evacuation. Non-nucleofected cells were taken after the washes in HBSS and seeded in 2ml of pre-cultured growth medium, serving as an additional control for FACS sorting.

#### 1.9 Selecting for single integration by FACS sorting

Nucleofected K562 cells were grown for 15 days to ensure that non-integrated plasmid DNA was eliminated, confirmed by the cells nucleofected without ZFNs. Sorting was performed with BD FACSAria II SORP (special-order research product). To sort cells that integrated the reporter construct successfully and in a single copy (∼4% of the population), a gate according to mCherry fluorescence was chosen so that only mCherry-expressing cells corresponding to a single copy of the construct were sorted (mCherry single integration population). The validity of this gate was verified by growing sorted cells for 8 additional days and re-examining mCherry levels, verifying that no cells exhibited mCherry levels corresponding to a double integration. A total of 7.5 million cells were collected in order to ensure adequate library representation. Master plasmid nucleofected cells were also sorted for single copy integration.

#### 1.10 Sorting single-integration library into 16 expression bins

Following single integration sorting, the mCherry single integration population was grown for 8 additional days before sorting into 16 bins according to the GFP/mCherry ratio. The bins were defined so they would span similar ranges of the ratio values, hence containing different percentage of the single integration population (from low expression to high - 2.5%, 4 bins of 8%, 9 bins of 6.5%, 5.5%, 1%). A total of 22 million cells were collected in order to ensure adequate library representation. The cells were grown further, and genomic DNA was purified from 5 million cells of each of the 16 bins, using DNeasy Blood & Tissue Kit (Qiagen) according to the manufacturer protocol.

#### 1.11 Preparing samples for sequencing

In order to maintain the complexity of the library amplified from gDNA, PCR reactions were carried out on gDNA amount calculated to contain an average of 200 copies of each oligo included in the sample. For each of the 16 bins, 20μg of gDNA were used as template in a two-step nested PCR in two tubes (to include the required amount of gDNA), each containing 100μl (in both steps); In the 1^st^ step each reaction contained 10μg gDNA, 50μl of Kapa Hifi ready mix X2 (KAPA biosystems), 5μl 10μM 5’ primer, and 5μl 10μM 3’ primer. The parameters for the first PCR were 95°C for 5 min, 18 cycles of 94°C for 30s, 65°C for 30s, and 72°C for 40s, each, and finally one cycle of 72°C for 5 min. Primers used for the first reaction were from the ActB promoter (5’-CTCTTCCTCAATCTCGCTCTCGCTC-3’) and the chimeric intron (5’-GACCAATAGGTGCCTATCAGAAACGC-3’). In the 2^nd^ PCR step each reaction contained 5μl of the first PCR product (uncleaned), 50μl of Kapa Hifi ready mix X2 (KAPA biosystems), 5μl 10μM 5’ primer, and 5μl 10μM 3’ primer. The PCR program was similar to the first step, using 24 cycles. For the second reaction the 5’ primer was comprised of a random 5nt sequence to increase complexity, followed by an 8nt barcode (one of three for each bin, underlined) and a library specific sequence (5’-HNHNH<u>XXXXXXXX</u>CGCGTCGCCGTGAGGAGG-3’). The common 3’ primer was (5’-HNHNHNHNGCCCCACGGAGGTGCCAC-3’. In both, the ‘N’s represent random nucleotides, and ‘H’ is A,C or T, in order to avoid synthesis of stretches of G that can affect initial clusters definition in NextSeq runs. The concentration of the PCR samples was measured using Quant-iT dsDNA assay kit (ThermoFisher) in a monochromator (Tecan i-control), and the samples were mixed in ratios corresponding to their ratio in the population. The library was separated from unspecific fragments by electrophoresis on a 2% agarose gel stained by EtBr, cut from the gel, and cleaned in 2 steps: gel extraction kit (QIAGEN), and SPRI beads (Agencourt AMPure XP). The sample was assessed for size and purity at the Tapestation, using high sensitivity D1K screenTape (Agilent Technologies, Santa Clara, California). 10ng library DNA were used for library preparation for NGS; specific Illumina adaptors were added, and DNA was amplified using 14 amplification cycles, protocol adopted from Blecher-Gonen et al(*60*). The sample was reanalyzed using Tapestation.

#### 1.12 Isolated clones measurements

Thirty isolated clones, at least one from each of the 16 expression bins were grown from single cells that were sorted into 96-wells plate. The clones were chosen based on their verified emergence from a single cell. After 28 days cell populations were analyzed in Flow Cytometry for eGFP expression and genomic DNA (gDNA) was purified. DreamTaq DNA polymerase (Thermo scientific) was used to amplify the library from 200ng gDNA, with same conditions and primers as in the library colony PCR. The PCR product was Sanger sequenced from the PCR Fw primer.

#### 1.13 Retroviruses production and infection

Phoenix virus packaging cells were used for retroviruses production as described before(*61*). 5*10^5^ cells were plated on 6cm plates 24hr prior to transfection. Cells were transfected with pPRIGp mChHA, a Moloney Murine Leukemia Virus (MMLV) retroviral plasmid expressing a bicistronic transcript encoding mCherry and eGFP separated by the EMCV Internal Ribosome Entry Site (IRES)(*57*). Each transfection included: 100μl DMEM with no serum or antibiotics, 12μl of FuGENE 6 transfection reagent (Promega) and 4μg of the retroviral plasmid. Transfection was performed according to the manufacturer’s instructions. After 24hr medium was replaced with fresh DMEM and H1299-EcoR cells were plated on 10cm plates for infection. After additional 24hr (48hr past transfection) 4ml of viruses-containing media were collected from Phoenix cells and centrifuged for 5 minutes in 1,500rpm. 3.5ml of viruses-containing media were added to 1.5ml RPMI media in each H1299 plate (total volume of 5ml) in addition to 5μl of Polybrene (AL-118, Sigma). After 24hr cells were washed 3 times with PBS, and fresh RPMI complete medium was added.

## 2. Data Analysis

### 2.1 Computing promoter activity threshold using empty vector measurements

In order to determine the activity threshold of core promoters we constructed and measured K562 cells for which we integrated an “empty vector” plasmids containing a linker sequence of 25bp between the AscI and RsrII restriction sites. We then measured the fluorophore levels of cells expressing the empty vector in flow cytometry using the same lasers intensities and settings as in the library sorting. We computed the normal distribution of the GFP/mCherry and extracted the mean and standard deviation (std). We set a threshold of 2 stds from the mean. Oligos with expression levels above this threshold (>=1.58) were considered as positive core promoters.

### 2.2 Mapping deep sequencing reads

DNA was sequenced on Illumina NextSeq-500 sequencer. To determine the identity of each oligo after sequencing we designed a unique 11-mer barcode upstream of the variable region. We obtained ∼42M reads for the entire library with a coverage of >=100 reads for 91% of the designed oligos (14,375 of 15,753). As reference sequence for mapping we constructed *in-silico* an “artificial library chromosome” by concatenating all the sequences of the 15,753 designed oligos with spacers of 50 ‘N’s. Single-end NextSeq reads in the length of 75 nucleotides, respectively, were trimmed to 45nt containing the common priming site and the unique oligo’s barcode. Trimmed reads were mapped to the artificial library chromosome using Novoalign aligner and the number of reads for each designed oligo was counted in each sample.

### 2.3 Computing mean expression and noise for each designed oligo

Deep sequencing reads from each bin were mapped using the unique 11bp barcode at the oligo 5’end. The distribution of reads across expression bins of each oligo was smoothed using the default ‘moving average’ method of MATLAB toolbox. Oligos with less than 100 reads after the smoothing process were filtered and ‘NaN’ values were assigned. Next, we detected the peak that contains the largest fraction of reads and that spans at least 3 adjacent bins. If obtained, additional smaller peaks were considered as technical noise as described before(*25*). We used the chosen peak to compute both mean expression and standard-deviation. Noise was quantified as the squared coefficient of variation (CV^2^), that is the variance divided by the square mean(*24*).

### 2.4 Statistical analysis

To assess the difference between expression levels of two groups that are distributed normally (e.g., native core promoters with and without TATA elements) we used a two-sample t-test. When expression levels were not distributed normally, such as in the case of the pre-initiation complex (PIC) binding sequences, we performed non-parametric Wilcoxon rank-sum test (for two samples) or Kruskal-Wallis test (for >2 samples). To compare the expression of pairs of sequences (e.g., adding a poly(dA:dT) tract to a sequence with two TF binding-sites) we performed Wilcoxon signed rank test. To evaluate differences in the variance resulting from the ZFN and retroviruses systems two-sample *F*-test for equal variances was performed. To examine significant differences between the proportions of positive core promoters in two groups (e.g., promoters and enhancers regions) we performed a two-proportion z-test. All correlations reported in the manuscript and the corresponding p-values were computed using Pearson correlation.

### 2.5 Fitting a logistic function

To examine the relationship between the number of binding-sites and promoter activity we fitted a logistic function with three parameters: maximal expression levels (*L*), the steepness of the curve (*k*), and the number of binding-sites at the sigmoid’s midpoint (*X*_*0*_).

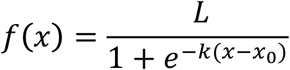

Oligos with expression levels below the activity threshold were filtered out. To test the agreement between the data and the fitted function we computed for each binding-site in each background the correlation and p-value between the measured expression levels and the fitted values.

